# Discovery, Characterization and Synthetic Potential of Two Novel Bacterial Alcohol Oxidases

**DOI:** 10.1101/2024.03.22.586125

**Authors:** Paula Cinca-Fernando, Christian Ascaso-Alegre, Emma Sevilla, Marta Martínez-Júlvez, Juan Mangas-Sánchez, Patricia Ferreira

## Abstract

The search for novel synthetic tools to prepare industrial chemicals in a safer and greener manner is a continuing challenge in synthetic chemistry. In this manuscript, we report the discovery, characterization, and synthetic potential of two novel aryl-alcohol oxidases from bacteria which are able to oxidize a variety of aliphatic and aromatic alcohols in high efficiencies (up to 4970 min^-1^mM^-1^). Crystal structures revealed unusually wide-open entrance to the active-site pockets compared to that previously described for traditional fungal aryl-alcohol oxidases, which could correlate with differences in substrate scope, catalytic efficiency, and other functional properties. Preparative-scale reactions and ability to operate at high substrate loadings also demonstrate the potential of these enzymes in synthetic chemistry with turnover numbers > 30000. Moreover, their availability as soluble and active recombinant proteins enabled their use as cell-free extracts which further highlights their potential for the large-scale production of carbonyl compounds.

## Introduction

The carbonyl moiety represents one of the most useful functional groups in synthetic chemistry and constitutes a synthetic crossroad from which a diverse array of functional groups can be accessed.[1] Particularly, aldehydes are the starting materials in many useful chemical asymmetric transformations such as Michael additions or the aldol and Mannich reactions.[2, 3] Furthermore, they are also present as the final compounds in many products from the food, pharma and cosmetic industry.[4, 5]. Traditionally, carbonyl groups are generated through alcohol oxidation, a process generally associated with hazardous conditions and the use of stoichiometric reagents.[6, 7] These conventional approaches often result in laborious downstream processes and lack regio- and chemo-selectivity, leading to the generation of by-products and requiring additional purification steps. In contrast, biocatalytic oxidations offer a greener approach to conventional strategies, as enzymes are highly efficient catalysts that display high selectivities and can function under mild reaction conditions, providing safer and environmentally friendlier processes.[8, 9] Amongst the different options, flavin-dependent alcohol oxidases (FAD-AOx EC 1.1.3.X), belonging to glucose-methanol-choline (GMC) superfamily of oxidoreductases, catalyse a broad variety of chemical transformations representing a promising and valuable tool in industrial biotechnology. For alcohol oxidation, FAD-AOx utilise the flavin cofactor as the hydride acceptor, which is regenerated upon reaction via aerobic oxidation, yielding H_2_O_2_ as the by-product. Therefore, oxygen is employed as the terminal oxidant and no external coenzymes are required in the process. Even though oxidases represent around 5-10% of the total genes in prokaryotic and eukaryotic genomes, the industrial application of this class of enzymes has been hampered by their limited availability.

Members of this superfamily include, among others, specific alcohol oxidases (EC. 1.1.3.13) acting on small aliphatic alcohols, and aryl alcohol oxidases (AAO, EC, 1.1.3.7) which can oxidise a broad range of activated primary alcohols (benzyl or allylic systems generally).[10] In the last decade, AAO have gained considerable interest as catalysts for aerobic alcohol oxidation due to their broad scope and the high efficiency displayed.[11, 12] Particularly, the AAO from *Pleurotus eryngii* (*Pe*AAO) and other fungi have been the focus of intensive study of their application in organic synthesis by several groups.[13-15] However, these enzymes require eukaryotic expression systems or tedious *in vitro* refolding procedures when expressed in *Escherichia coli* to produce active catalysts, which limits their academic and industrial uptake for general alcohol oxidation. In efforts to overcome these limitations, several enzymes such as a choline oxidase from *Arthrobacter chlorophenolicus* and a methanol oxidase from *Phanerochaete chrysosporium* have been evolved to obtain enhanced variants able to oxidise typical AAO alcohol substrates.[16, 17]

Given the widespread natural occurrence of FAD-AOx across all kingdoms, we hypothesise that bacteria could represent a promising reservoir to discover new AAOs with an expanded synthetic scope. Moreover, enzymes from bacterial sources are frequently easily expressed in *E. coli*, which is an ideal host for recombinant protein expression in biocatalysis due to its rapid growth, easy genetic manipulation and straightforward purification protocols. In this study, we present our findings on the discovery, characterisation, and evaluation of the synthetic potential of two novel AAO originating from *Sphingobacterium daejeonense* (*Sd*AAO) and *Streptomyces hiroshimensis* (*Sh*AAO).

## Results and Discussion

### Identification and selection of new bacterial AAO

In the search for new prokaryotic biocatalysts to produce aliphatic and aromatic aldehydes from the corresponding primary alcohols, we blast the amino acid sequence of *Pe*AAO in bacterial NCBI database. A total of 12 sequences were identified in *Actinomadura, Geminicoccaceae, Streptomyces, Sphingobacterium* genera (**Table S1**). These putative AAO sequences contain the highly conserved histidines in the adjacent motif to the C-terminal as well as the ADP-binding domain and consensus PS00623 and PS00624 sequences characteristic of GMC proteins (**Figure 1**). Moreover, the identified sequences form a distinct subgroup closely related to those fungal sequences in AAO-GMC clade, clearly separated from the MOX-GMC group in the maximum likelihood tree (**Figure S1)**. The molecular models of the identified proteins, generated from Alphafold, predict similar fold topology to other known members of the GMC superfamily (**Figure S2A**). Notably, a detailed examination of bacterial putative substrate binding domains revealed differences in amino acid composition and accessibility of their access channels (**Figure S2B-C**). These differences may potentially result in a broader variety of substrate specificities. Thus, one representative sequence from each identified specie was selected for overproduction as recombinant protein in *E. coli* (highlighted sequences in **Table S1 and S2**).

**Figure 1.**
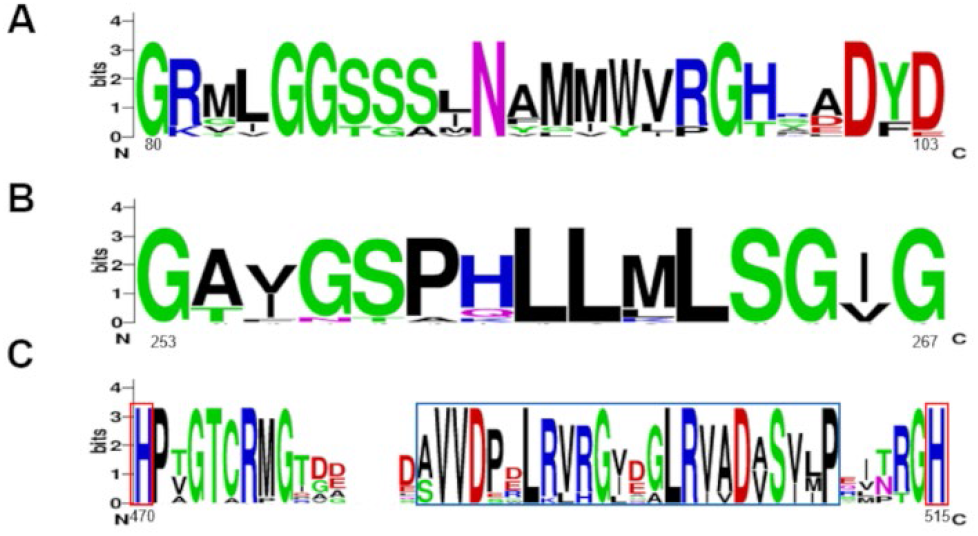
Sequence logo for 12 identified proteins corresponding to the consensus PS00623 (A) and PS00624 (B) sequences and the C-terminal adjacent motif bordered in blue (C). The conserved histidine residues bordered in red. The numbers indicate the position in the *Sd*AAO sequence and the heights of letters in each column indicate the relative frequency of each amino acid. Representation generated using Weblogo.[18]

### Expression and purification of two novel bacterial AAO

The nine optimised genes were subcloned into pET28a vector and transformed into *E. coli* C41 (DE3) strain. To optimise its expression as soluble proteins, transformed cells were culture simultaneously in autoinduction and IPTG-induced LB at 20 and 37 ºC for 4-72 hours (post-induction in the case of LB). For *Sd*AAO and *Sh*AAO, the highest levels of soluble proteins were achieved with LB at 37 ºC for 48 and 24 hours, respectively, making these conditions optimal for large-scale protein production. However, for the remaining proteins, analysis of cell extracts revealed its accumulation in the insoluble fraction, even when using the *E. coli* Rosetta (DE3) strain as alternative expression host. Therefore, *Sd*AAO and *Sh*AAO could be purified in a single affinity chromatographic step, with yields over 20 mg of protein per L of culture. SDS-PAGE analysis revealed a single band of about 55 kDa, in good agreement with the theoretical molecular weights of 59 and 55 KDa for *Sd*AAO and *Sh*AAO, respectively (**Figure S3A-B**). The *Sd*AAO and *Sh*AAO spectra exhibited the typical bands I and II of the flavin, indicating that the flavin cofactor was in the oxidised state and correctly incorporated into the proteins (**Figure S3C** and **Table S3**). Moreover, *Sd*AAO and *Sh*AAO eluted in gel filtration chromatography as a single peak, displaying apparent molecular weights of 50 and 41 kDa, respectively, corresponding to their respective monomeric forms (data not shown).

### Operational stability, substrate scope and kinetic properties

The operational window regarding temperature and pH of the two novel AAOs was initially evaluated. *Sh*AAO and *Sd*AAO showed similar thermoestability with thermal melting temperature (*T*_m_) values of 50.9 and 48.6 ºC, respectively (**Figure 2A**). We also evaluated the thermostability by measuring the residual activity of both enzymes towards the oxidation of 2,4-hexadien-1-ol after incubation at different temperatures. As expected, *Sh*AAO and *Sd*AAO kept 50% of their activity after 10-min incubation at 47.8 and 46.8 °C (*T*_50_^10^ values), respectively (**Figure 2B**). In terms of pH stability over time, *Sd*AAO exhibited robust performance, retaining over 80% of its initial activity after 72 h of incubation in the pH range of 5 to 7, and maintaining 70% at pH 8-9 (**Figure 3A**). In contrast, *Sh*AAO showed a more limited pH stability profile, with over 80% remaining activity only at pH 6 (**Figure 3B**). Its activity declined rapidly at other pH values, with complete loss observed within 24 to 48 hours. Both enzymes precipitated at pH 4, showing no activity at lower pH values. Additionally, no effect on *Sd*AAO stability was observed after 24 hours of exposure up to 26 mM of H_2_O_2_, maintaining even 70% of its initial activity at 35 mM (**Figure 3C**). However, *Sh*AAO showed decreased stability in the presence of H_2_O_2_ with its residual activity dropping to 10% after 24 hours incubation with 9 mM H_2_O_2_ (**Figure 3D**). The enzyme became inactivated at higher H_2_O_2_ concentrations, maintaining only 10% of initial activity after 11 hours of exposure with 35 mM.

**Figure 2.**
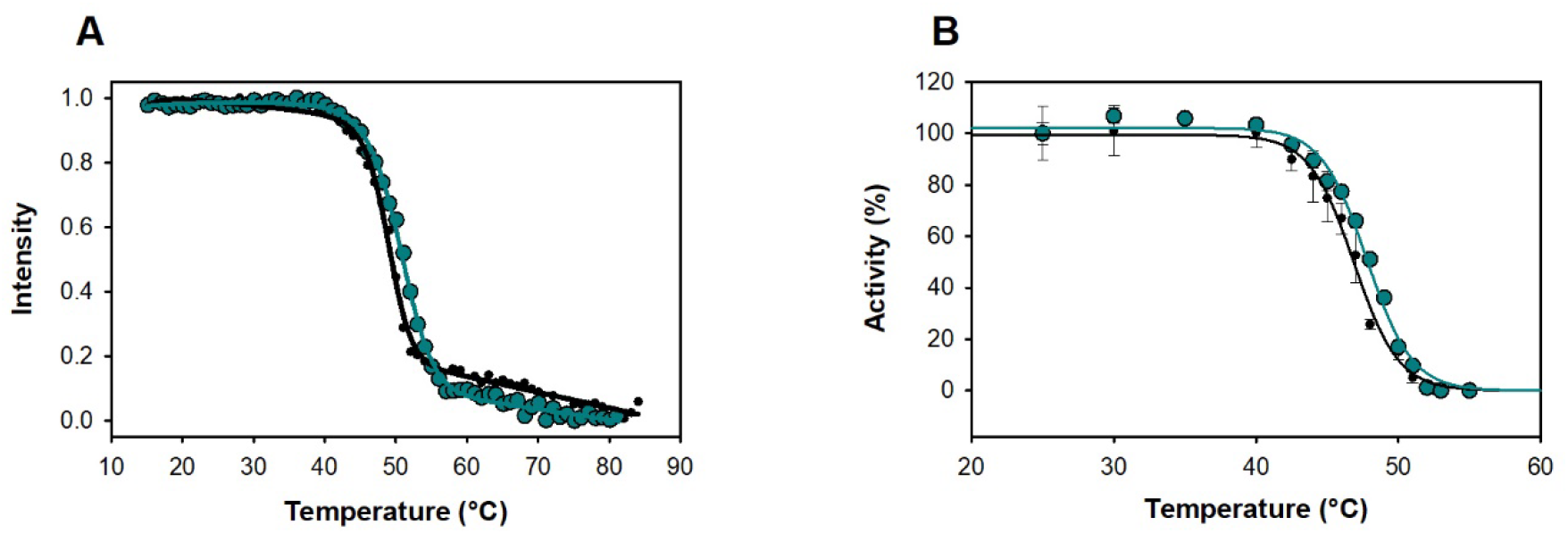
Influence of temperature on *Sd*AAO (black circles) and *Sh*AAO (cyan circles) stability. (A) Melting temperatures measured by FAD release upon protein denaturation. (B) Thermal stability estimated from residual activity after 10 minutes of incubation at different temperatures. Residual activity is given in %.

**Figure 3.**
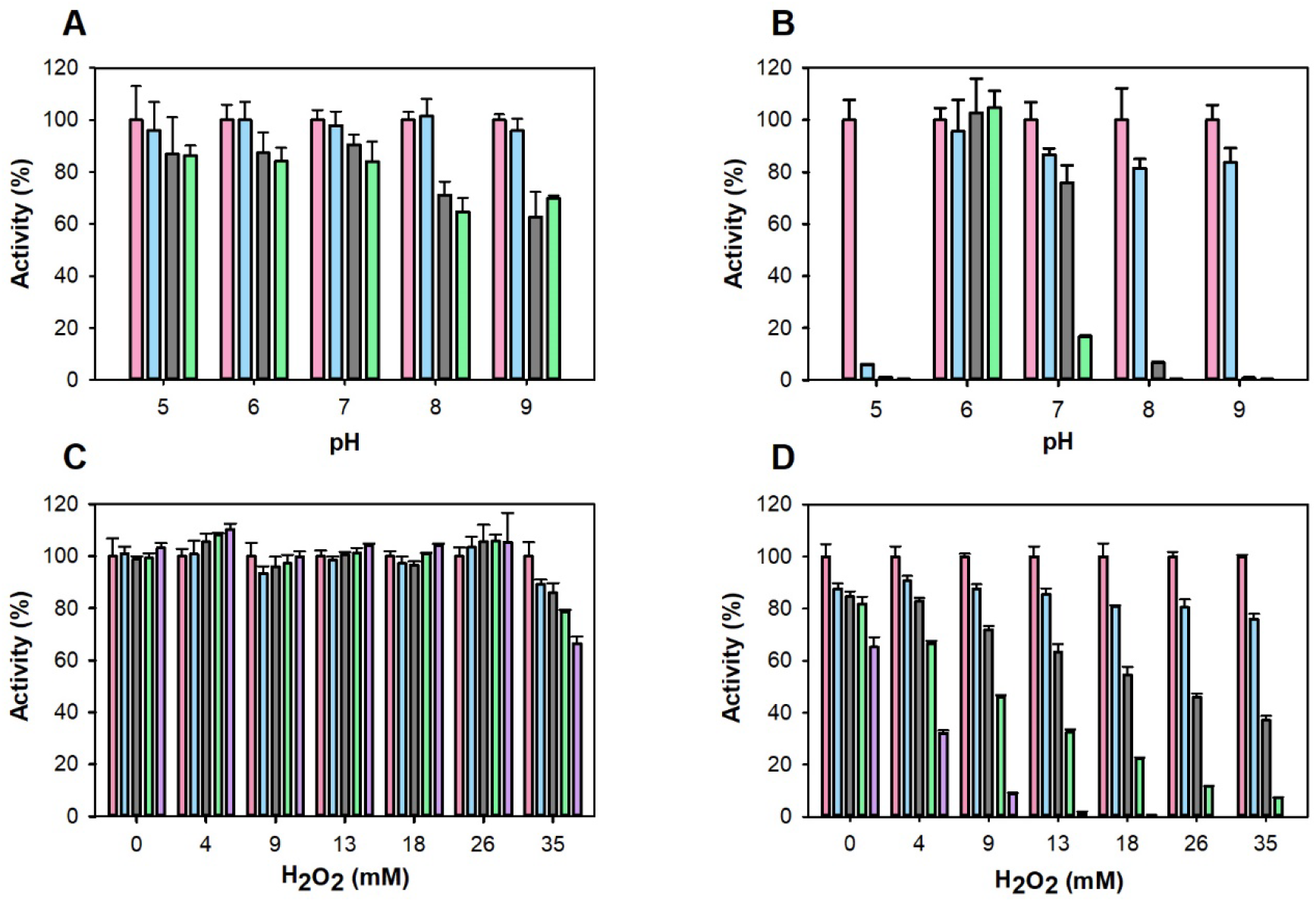
Effect of pH and H_2_O_2_ on *Sd*AAO and *Sh*AAO stability. pH stability of *Sd*AAO (A) and *Sh*AAO (B) was determined during 72-h incubation at 25°C in B&R buffer (pH 5 to 9). Activity was measured 5 minutes after mixing (pink), after 24 hours (blue), 48 hours (gray) and 72 hours (green). H_2_O_2_ stability of *Sd*AAO (C) and *Sh*AAO (D) was determined during 24-h incubation at 25 °C in Tris 50 mM pH 7. Activity was measured 5 minutes after mixing (pink), after 3 hours (blue), 6 hours (gray), 11 hours (green) and 24 hours (purple). Residual activity is given in %.

The pH optimum of *Sh*AAO and *Sd*AAO was determined in the pH range of 3 to 12 using the oxidation of 2,4-hexadien-1-ol (**Figure S4**). Notably, *Sh*AAO displayed a broader optimal pH range (pH 7-10) compared to *Sd*AAO (pH 6-7). Specifically, *Sh*AAO retained ∼ 90% of its activity over a pH range from 5 to 11. In contrast, *Sd*AAO activity decayed below 70% at both one pH unit below and above its optimum, highlighting a narrower pH tolerance.

A FOX screening method based on the Fenton reaction, wherein the H_2_O_2_ byproduct of alcohol oxidase activity is quantified in a highly sensitive manner, was initially used to assess the substrate scope of *Sh*AAO and *Sd*AAO. A total of 15 compounds spanning benzylic, other cyclic, and aliphatic primary alcohols as well as secondary alcohols were tested (**Figure S5** and **Table S4**). For comparative description of substrate specificity, the quantified activity for each enzyme towards 2-4-hexadien-1-ol was standardized to 100%. *Sh*AAO exhibited the highest relative activities towards aliphatic-polyunsaturated primary alcohols with conjugated double bonds, 2,4-hexadien-1-ol (100%) and *3,7-* dimethyl-2,6-octadien-1 *(92%)*, followed by 3-phenyl-2-propen-1-ol alcohol (cinnamyl alcohol, 88%). In contrast, the rest of benzyl alcohols, low (<5%) or no activity was detected, in opposition to substrate scopes previously described for fungal AAOs.[11, 19, 20]

On the other hand, *Sd*AAO preferentially oxidized aromatic primary alcohols, showing the highest activities with cinnamyl, 4-nitrobenzyl and 4-bromobenzyl alcohols (390%, 255% and 119%, respectively) which is in accordance to the scope reported for fungal AAOs.[11, 19, 20] In both enzymes, benzylic substrates presenting an extended unsaturated side chain, as in cinnamyl alcohol, resulted in increased activity compared to 3,4-dimethoxybenzyl alcohol (5- and 88-fold increase for *Sd*AAO and *Sh*AAO, respectively) in agreement to that described for fungal AAOs. In the case of *Sd*AAO, the presence of electron-withdrawing groups on 4-bromobenzyl and 4-nitrobenzyl alcohols had a positive effect, leading to 1.5- and 3.5-times higher relative activity than that observed for substrates bearing electron-deficient aryl groups like 3,4-dimethoxybenzyl alcohol. Lower activities were also observed in the oxidation of heteroaryl alcohols such as 2-Pyridinemethanol and 3-(hydroxymethyl)pyridine (35% and 1%, respectively). Notably, 5-(hydroxymethyl)furfural (HMF), a compound of interest for its use in the synthesis of bioplastics, was readily oxidized by both enzymes (with 69% and 18% relative activity for *Sd*AAO and *Sh*AAO, respectively) consistent with previous reports for fungal AAOs.[19, 21] Regarding secondary alcohols, both enzymes were able to oxidase aromatic substrates 4-phenyl-2-butanol and 1-phenylethanol, with relative activities comparable to that observed for heteroaryl primary alcohol 2-pyridinemethanol.

Steady-state parameters were determined for oxidation of a representative sample of primary alcohols based on the substrate profiling (**Table 1 and Figure S6**). *Sd*AAO exhibited broader substrate specificity compared to *Sh*AAO, enabling the determination of kinetic parameters for all tested alcohols. However, *Sh*AAO showed the highest turnover number with 2,4-hexadien-1-ol followed by cinnamyl alcohol and 3,7-dimethyl-2,6-octadien-1-ol (*k*_cat_ values of 891, 173 and 110 min^-1^, respectively). A significant decrease in oxidation rates towards 4-bromobenzyl and 4-nitrobenzyl alcohols were observed, with *k*_cat_ values of 12 and 2 min^-1^, respectively. Additionally, *Sh*AAO exhibited moderate substrate inhibition with 4-nitrobenzyl alcohol, with a constant inhibition (*K*_i_) value of 19 μM (9-fold higher than the corresponding *K*_m_ value). Concerning *Sd*AAO, *k*_*cat*_ values up to 54 min^-1^ were found for benzylic alcohols. Notably, *Sd*AAO also displayed greater affinity (up to a 326-fold lower *K*_M_) than *Sh*AAO, which resulted in higher catalytic efficiencies for *Sd*AAO across the panel, except for 2,4-hexadien-1-ol, ranging from 36 to 350-fold higher efficiencies compared to those obtained for *Sh*AAO. In general terms, *Sd*AAO and *Sh*AAO exhibited lower turnovers than those previously described for fungal ones, but similar or even higher affinities for bulky substrates such as cinnamyl alcohol.[11, 20, 22]

**Table 1.**
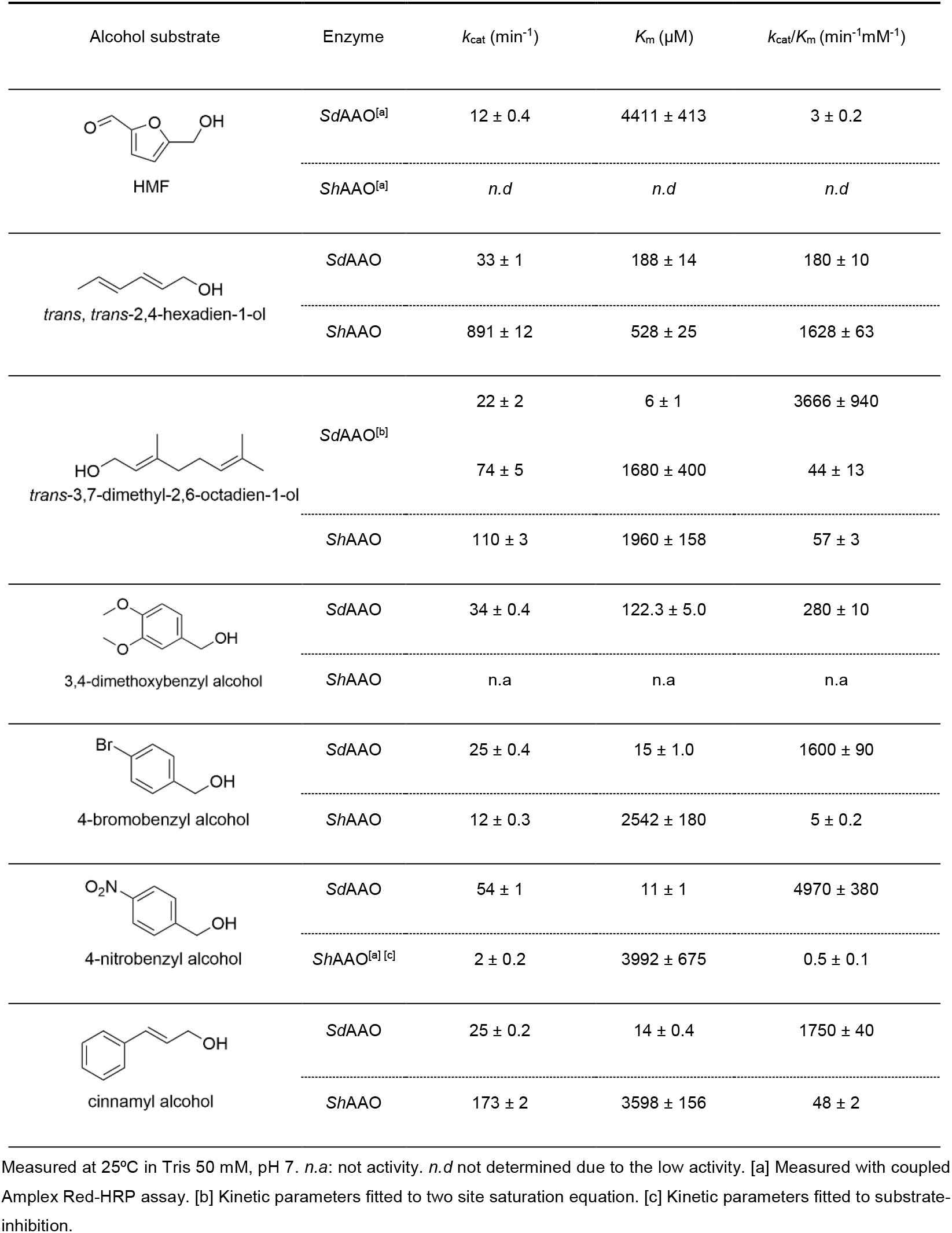
Steady-state kinetic parameters of *Sd*AAO and *Sh*AAO for the oxidation of different alcohols.

| Alcohol substrate | Enzyme | $k_{cat}$ (min<sup>-1</sup>) | $K_m$ (μM) | $k_{cat}/K_m$ (min<sup>-1</sup>mM<sup>-1</sup>) |
| --- | --- | --- | --- | --- |
| HMF | *SdAAO*<sup>[a]</sup> | 12 ± 0.4 | 4411 ± 413 | 3 ± 0.2 |
|  | *ShAAO*<sup>[a]</sup> | *n.d* | *n.d* | *n.d* |
| *trans*, *trans*-2,4-hexadien-1-ol | *SdAAO* | 33 ± 1 | 188 ± 14 | 180 ± 10 |
|  | *ShAAO* | 891 ± 12 | 528 ± 25 | 1628 ± 63 |
| *trans*-3,7-dimethyl-2,6-octadien-1-ol | *SdAAO*<sup>[b]</sup> | 22 ± 2 | 6 ± 1 | 3666 ± 940 |
|  |  | 74 ± 5 | 1680 ± 400 | 44 ± 13 |
|  | *ShAAO* | 110 ± 3 | 1960 ± 158 | 57 ± 3 |
| 3,4-dimethoxybenzyl alcohol | *SdAAO* | 34 ± 0.4 | 122.3 ± 5.0 | 280 ± 10 |
|  | *ShAAO* | *n.a* | *n.a* | *n.a* |
| 4-bromobenzyl alcohol | *SdAAO* | 25 ± 0.4 | 15 ± 1.0 | 1600 ± 90 |
|  | *ShAAO* | 12 ± 0.3 | 2542 ± 180 | 5 ± 0.2 |
| 4-nitrobenzyl alcohol | *SdAAO* | 54 ± 1 | 11 ± 1 | 4970 ± 380 |
|  | *ShAAO*<sup>[a] [c]</sup> | 2 ± 0.2 | 3992 ± 675 | 0.5 ± 0.1 |
| cinnamyl alcohol | *SdAAO* | 25 ± 0.2 | 14 ± 0.4 | 1750 ± 40 |
|  | *ShAAO* | 173 ± 2 | 3598 ± 156 | 48 ± 2 |

Measured at 25°C in Tris 50 mM, pH 7. *n.a*: not activity. *n.d* not determined due to the low activity. [a] Measured with coupled Amplex Red-HRP assay. [b] Kinetic parameters fitted to two site saturation equation. [c] Kinetic parameters fitted to substrate-inhibition.

### Structural features of bacterial AAOs with major accessibility to active sites

The two bacterial crystal structures were solved at a resolution of 2.0 Å, and each chain (A and B in *Sd*AAO and A in *Sh*AAO) show one FAD molecule and one alcohol molecule in the active pocket (a hexane-1,6-diol (HEZ) molecule in *Sd*AAO, and a triethylene glycol (PGE) molecule in *Sh*AAO) derived from the corresponding crystallization conditions. Between them, the overall structural comparison indicates a high folding similarity with an RMSD value of 0.85 Å (for 354 Cα atoms superimposed). This fold topology resembles that of other members of GMC superfamily such as *Pe*AAO, characterised by two major domains (**Figure 4B**), resulting in RMSD values of 1.41 Å and 0.87 Å for the Cα atoms and sequence identities of 34.7 % and 37.5 % in *Sd*AAO and *Sh*AAO, respectively. In both proteins, the FAD-binding domain consists of a five-stranded parallel β-sheet interrupted by a three-stranded antiparallel β-sheet that connects with two antiparallel β-strands, along with seven α-helices. The FAD cofactor is noncovalently bound to both proteins via hydrogen bounds to surrounding residues and several water molecules (**Figure S7**). The substrate-binding domain in both *Sd*AAO and *Sh*AAO contains six and five antiparallel β strands, respectively, flanked by three α-helices. Additionally, another helix is located on the opposite side of the domain (315-324 *Sd*AAO and 311-320 *Sh*AAO). Interestingly, both structures have a substrate analogue molecule bound in the active-site cavity, with the site being highly accessible (**Figure 5**).

**Figure 4.**
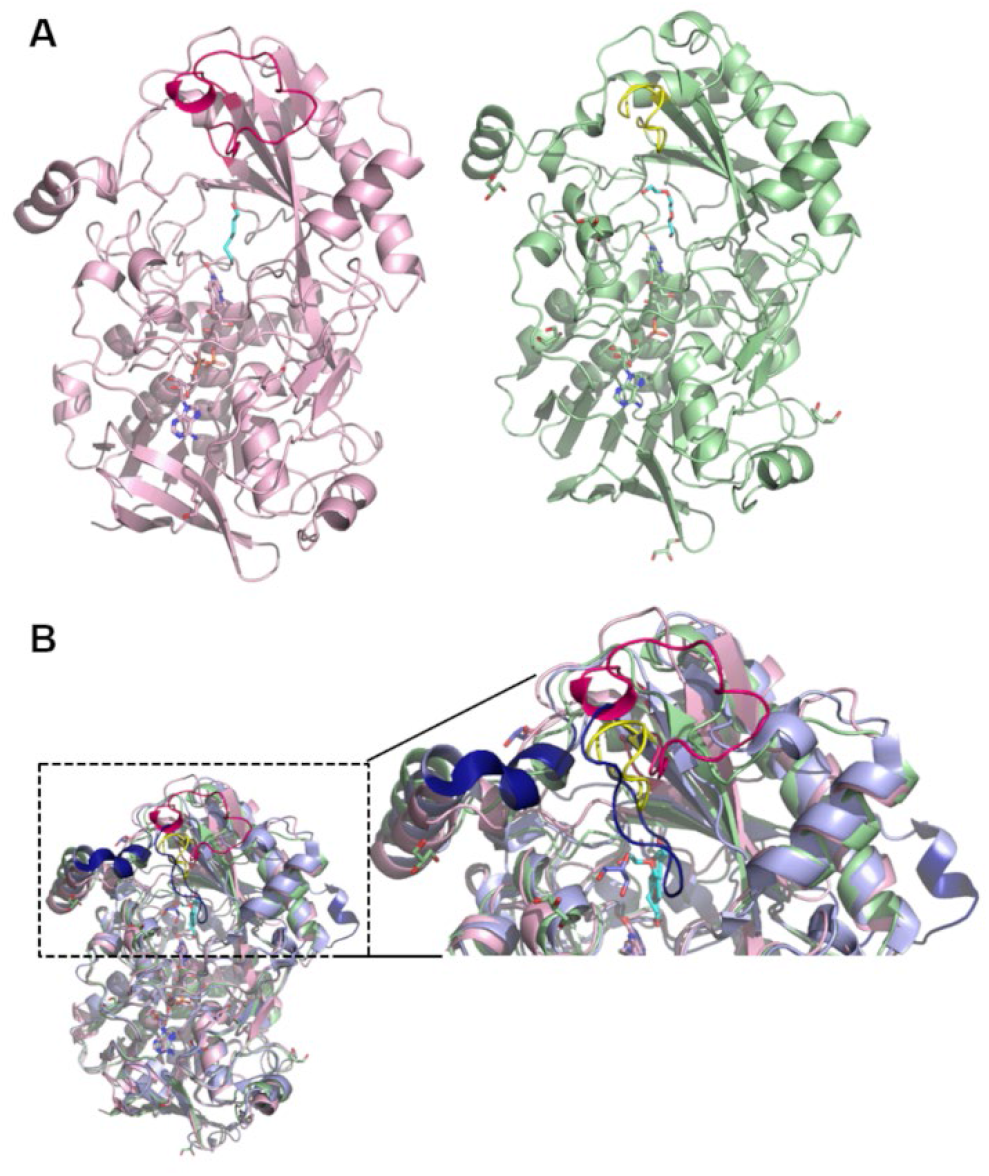
The crystal structures in cartoon representation of (A) *Sd*AAO (light pink, PDB ID: 8RPF) and *Sh*AAO (light green, PDB ID: 8RPG) and (B) a superposition of the bacterial AAOs with *Pe*AAO (light blue, PDBID: 5OC1). Besides, an enlarged image showing the main structural differences among these structures (highlighted in dark blue in PeAAO). The alcohol molecules in bacterial AAOs and the 4-methoxybenzoic acid in *Pe*AAO are displayed with cyan carbons. Loops that seem to modulate the opening of the channel to the active site in each structure are colored in hotpink, yellow and dark blue for *Sd*AAO, *Sh*AAO and *Pe*AAO, respectively.

**Figure 5.**
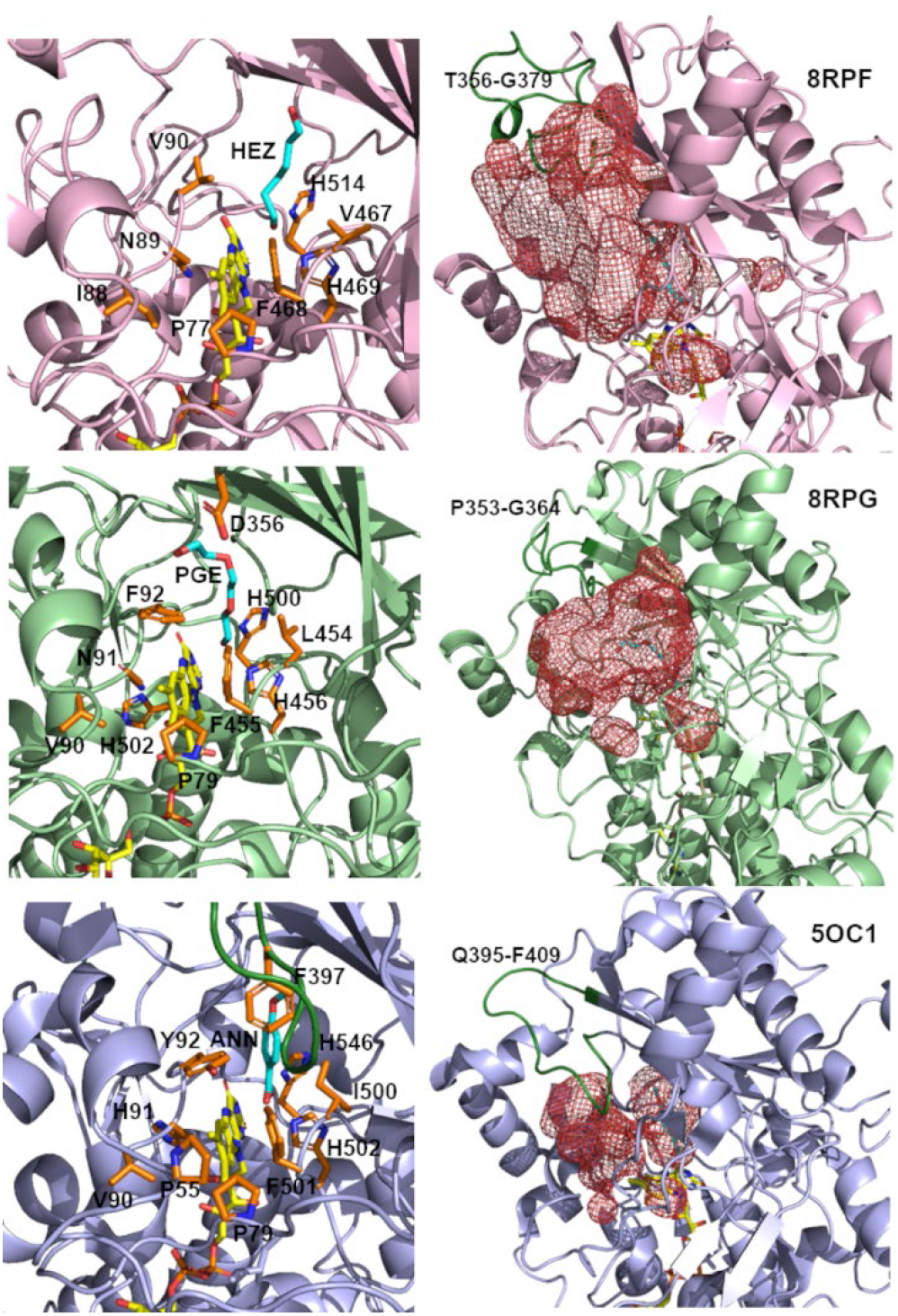
Substrate binding sites and access channels to the active sites in *Sd*AAO (pink), *Sh*AAO (green) and *Pe*AAO (blue) complexes. In all structures, residues lining the pathway to the active site and key residues in the active site are shown as sticks with carbon atoms in orange. HEZ, PGE and ANN are displayed with cyan carbons. The access channels and active-site pockets are shown as red meshes in the corresponding right-hand images (red, calculated by HOLLOW; Ho & Gruswitz, 2008).[26] The loop restricting access to the channel in *Pe*AAO and the corresponding ones in the bacterial structures are coloured dark green.

This internal cavity is located in front of the *re* side of the FAD isoalloxazine and flanked by V90, F468, His469 and His514 in *Sd*AAO, and by F92, F455, His456 and His500 in *Sh*AAO, similarly to that described for *Pe*AAO (**Figure 5**). Notably, HEZ and PGE are hydrogen-bonded to the N5 atom of FAD isoalloxazine ring via their O1 atoms, as well as to the NE2 and ND1 atoms of His469 and His501 in *Sd*AAO, and His456 and His500 in *Sh*AAO. The position of His469 in *Sd*AAO and His456 in *Sh*AAO corresponds to the highly conserved active site histidine residue that activates alcohol substrate by proton abstraction during the reductive half-reaction in the majority of GMC oxidoreductases[23, 24]. Thus, the substrate-like binding mode is compatible with redox catalysis and suggests that *Sd*AAO and *Sh*AAO may share the consensus hydride transfer mechanism, assisted by a catalytic base, as previously described most well-known characterized GMC members. Despite similarities among bacterial and fungal AAOs structures, significant differences primarily exist in three structural components (**Figure 4B** and **Figure 5**). *Pe*AAO has a helical insertion that is lacking in bacterial AAOs, broadening its structure (**Figure 4B**). However, the most significant difference is in two structural elements in the surroundings of their active sites that modulate the entrance of substrates in the fungal AAO.[25]

In *Pe*AAO, a α-helix compressing residues 326-334 (**Figure 4B**) and a 14-residue loop containing F397 (compressing residues 395-409, **Figure 5**) between the two antiparallel β strands of the substrate-binding domain, restrict access to active site through a hydrophobic bottleneck (**Figure 5**).[27] In contrast, bacterial structures lack this protruding α-helix and the equivalent loops (compressing residues 353-364 and 356-379 in *Sh*AAO and *Sd*AAO, respectively) adopt a conformation that do not cover the entrance to the channel in the same extent as *Pe*AAO does, creating wide and unblocked active sites, especially in *Sd*AAO structure (**Figure 5**). Thus, the surface areas covering the active-site pocket were established to be approximately 2026 Å^2^ for *Sd*AAO and 1383 Å^2^ *Sh*AAO considerably higher than the 903 Å^2^ estimated for *Pe*AAO. Interestingly, the fully accessible catalytic tunnel of *Sd*AAO is similar to that of previously described for the fungal AAO from *Thermothelomyces thermophilus*, despite significant differences in the general folding of both proteins, particularly in the substrate-binding domain which might responsible of its lower catalytic efficiency.[28] These differences in substrate accessibility to the active site might explain the observed differences in substrate specificities among proteins. Such insights are of interest for expanding the typical substrate scope of AAO proteins to include bulky substrates.

### Biotransformations

Due to the high turnover rates observed in steady-state kinetic studies as well as the distinct scope displayed compared to fungal AAOs, we selected *Sh*AAO for an assessment of the potential use of these enzymes in synthetic chemistry. We initially performed a series of analytical scale biotransformations with the oxidation of cinnamyl alcohol serving as the model reaction using 8.8 μM *Sh*AAO in pH 7 100 mM Tris-HCl buffer at 30 ºC. After 24h, 56% conversion to cinnamaldehyde was obtained at 20 mM substrate concentration (**Table 2** and **Figures S8 and S9**). The addition of bovine liver catalase (2-5 kU) to remove H_2_O_2_, resulted in a notable increase in conversion (up to 96%) so we continued the evaluation at higher substrate concentrations (**Table 2, entries 2-6**). Remarkably, the enzyme exhibited tolerance up to an 80 mM substrate concentration without showing any inhibition issues. Following optimisation of buffer, pH, catalase and enzyme concentrations, a remarkable TON of 31290 was achieved at 40 mM substrate concentration (**entry 13**, NaPi buffer pH 6, 30 °C, 1.1 μM *Sh*AAO, catalase added in two portions), with a turnover frequency (TOF) of 309 min^-1^ under the optimised conditions (**Figure S8**). High levels of protein expression in *E. coli* offers the prospect of using semi purified enzyme preparations such as cell-free extracts (CFE).

**Table 2.**
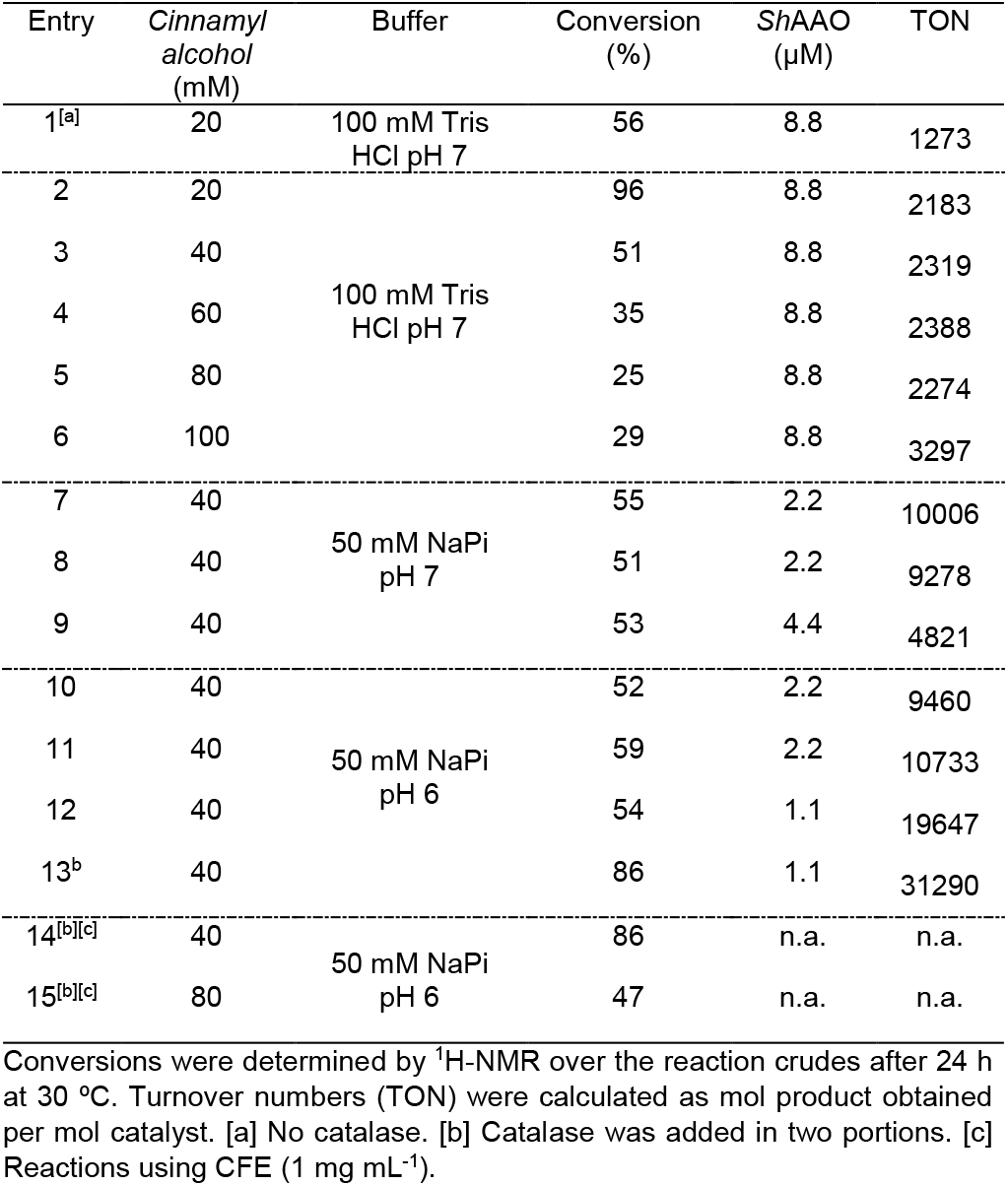
Reaction optimization of analytical-scale biotransformations.

These formulations are a preferred option in industry due to their economic advantages, as it avoids the need for supplementary protein purification steps.[29] Under optimised conditions and using 1 mg mL^-1^ CFE, 86% conversion to cinnamaldehyde were obtained at 40 mM substrate concentration (**Table 2, entry 14**). To further evaluate the synthetic applicability of *Sh*AAO, a preparative-scale biotransformation was performed starting from 1 mmol (134 mg) of cinnamyl alcohol at a 40 mM concentration (5.4 g L^-1^) using purified *Sh*AAO added in two portions (5.25 mg total enzyme amount, 0.21 g L^-1^). The reaction progress was monitored by thin-layer chromatography and subsequently purified by extraction and flash chromatography, obtaining cinnamaldehyde in an 83% isolated yield. This corresponds to a remarkable TON of 9024 and a catalyst productivity of 21 g product/g enzyme.

## Conclusions

Recent studies have shown the tremendous potential of oxidases in synthetic chemistry due to their exquisite selectivity and mild reaction conditions that generally leads to safer procedures with a lower environmental toll.[29],[30] In summary, we report the discovery and characterisation of two novel bacterial AAOs from *Sphingobacterium daejeonense* (*Sd*AAO) and *Streptomyces hiroshimensis* (*Sh*AAO) which have been shown to possess promising potential for their use in synthetic chemistry as alcohol oxidation catalysts under mild reaction conditions. While *Sd*AAO demonstrated a scope similar to that of previously reported fungal AAOs, *Sh*AAO exhibited distinctive substrate tolerance, displaying a preference for long-chain aliphatic and aromatic allylic alcohols. Analysis of crystal structures unveiled unusually wide accessible catalytic tunnels, contrasting with those previously described for traditional fungal AAOs, which may underlie variations in substrate specificity and additional functional properties. This divergence provides valuable insights for potential modifications through protein engineering, aiming to produce variants with expanded scopes and enhanced efficiencies that will contribute to extending the applications of these enzymes. We further demonstrated their synthetic potential by performing reactions at high substrate concentrations as well as preparative-scale reactions and proven the suitability of using semi-purified enzyme preparations. Finally, this study evidences the natural distribution of aryl-alcohol oxidases in bacteria, opening paths for the discovery of new biocatalysts to produce valuable aldehydes.

## Materials and Methods

### Screening for bacterial AAO genes and sequence analysis

The search of novel AAO-like sequences was performed by querying bacterial non-redundant protein database from the National Center for Biotechnology Information (NCBI) with a fungal AAO from *P. eryngii* (NCBI code AAC72747.1) as reference, and employing the protein Basic Local Alignment Search Tool (BLAST). Multiple alignments of sequences exhibiting the highest similarities were conducted with MUSCLE to identify the conserved motifs, including ADP binding domain and Prosite PS00623 and PS00624 sequences, as well as highly conserved histidine residues in GMC proteins.[10, 31] Following sequence analysis, a maximum-likelihood phylogenetic tree was constructed using MEGA version 11.0.13 with 1000-iteration bootstrapping using the Whelan and Goldman model of evolution with gamma-distributed rate variation with empirical amino acid frequencies and invariant sites (WAG+F+I+G).[32] Furthermore, 3D structural model of the selected proteins was predicted using AlphaFold server.[33]

### Production of recombinant proteins

The codifying DNA sequences of selected proteins (Table S1) were heterologously expressed in *E. coli* as recombinant proteins with an N-terminal His_6_-tag using the pET-28a(+) vector. Proteins were expressed and purified as described in the supplementary materials.

### Spectroscopic studies

UV-visible spectra of purified AAOs were measured between 250 and 800 nm. Protein concentrations were determined using their molar absorption coefficients, estimated by denaturation the protein with 3M guanidinium chloride in Tris-HCl 50 mM pH 7 (standard buffer), followed by quantification of the released FAD (ε_450_ = 12250 M^-1^ cm^−1^). [34] The extinction coefficients for *Sd*AAO and *Sh*AAO were ε_453nm_ = 11488 M^-1^ cm^−1^ and ε_456nm_ = 8929 M^-1^ cm^−1^, respectively.

### Substrate spectrum

To determine the substrate specificities of AAOs, a high-throughput screening was performed using the FOX method. This assay detects the production of H_2_O_2_ by coupling the Fenton reaction to xylenol orange forming a blue-purple complex, measurable at 560 nm (ε_560_ = 225000 M^-1^cm^-1^). Further details can be found in supporting information.

### Steady-state kinetics

Steady-state kinetic studies were conducted for selected alcohols at varying concentrations by monitoring their rate of oxidations to the corresponding aldehydes in standard buffer at 25 ºC. The measurements with HMF and 4-nitrobenzyl alcohol were performed by following the production of H_2_O_2_. See supporting information for more details.

### Operational stability assays pH and H_**2**_**O**_**2**_ **stability**

The pH and H_2_O_2_ stability as well as optimal pH were assessed by incubating the enzymes in 100 mM Britton and Robinson (B&R) buffer at different pHs ranging from 3 to 9 or in the presence of H_2_O_2_ (0 to 35 mM) in standard buffer at 25 °C. Residual activities were determined by monitoring the oxidation of 4 mM 2,4-hexadien-1-ol in standard buffer, at 25 °C. The melting temperatures of AAOs was determined by changes in the FAD fluorescence emission due to unfolding proteins. For determination of T_50_^10^, the temperature at which 50% of activity is retained after 10-min heat treatment, proteins were incubated in standard buffer, spanning temperatures from 25 to 60 °C for 10 min. See supporting information for more details.

### Crystallization, data collection and model resolution of *Sd*AAO and *Sh*AAO

For the crystallization of both proteins, several crystallization conditions were screenings using the sitting-drop vapour diffusion technique. Diffraction data were collected using the BL13-XALOC beamline of ALBA, Barcelona (Spain). Diffraction data sets were scaled using XDS and SCALA from CCP4i.[35, 36] The phasing step was performed by MOLREP from CCP4i using AlphaFold models of both proteins. [33, 37] Structure refinements were performed by alternating manual and automatic cycles using COOT and REFMAC5 of the CCP4 package.[38, 39] Data collection statistics are summarized in **Table S5**. The atomic coordinates have been deposited in the Protein Data Bank (PDB) with accession numbers 8RPF (*Sd*AAO) and 8RPG (*Sh*AAO).

## Acknowledgements

We would like to thank the Agencia Estatal de Investigación (AEI), the Ministerio de Ciencia y Universidades (Ministry of Science and Universities, MICIU), and the EU for the financial support (TED2021-130803B-I00 MICIU/AEI /10.13039/501100011033 NextGenerationEU/PRTR; PID2022-136369NB-I00 funded by MCIN/ AEI/10.13039/501100011033 and FEDER). J.M-S also thanks the AEI for a Ramón y Cajal Fellowship (RYC2021-032021-I). Authors would like to acknowledge the use of Servicio General de Apoyo a la Investigación-SAI, Universidad de Zaragoza.

## Notes

### Competing Interest Statement

The authors have declared no competing interest.

